# Arctic Ocean virus communities: seasonality, bipolarity, and prokaryotic interactions

**DOI:** 10.1101/2024.10.11.617772

**Authors:** Alyzza Calayag, Taylor Priest, Ellen Oldenburg, Jan Muschiol, Ovidiu Popa, Matthias Wietz, David M. Needham

**Author notes:** Institute for Chemistry and Biology of the Marine Environment, 26129 Oldenburg, Germany.

## Abstract

Viruses play important roles in ocean environments as agents of mortality and genetic transfer, influencing ecology, evolution and biogeochemical processes. However, we know little about the diversity, seasonality, and host interactions of viruses in polar waters. To address this, we studied dsDNA viruses in the Arctic Fram Strait across four years via 47 long-read metagenomes of the cellular size-fraction. Among 5,662 vOTUs, 69%, 30% and 1.4% were bacteriophages (Myoviridae, Podoviridae, and Siphoviridae), Unassigned, and Phycodnaviridae, respectively. Viral coverage was, on average, 5-fold higher than prokaryotic coverage, and 8-fold higher in summer. Viral community composition showed annual peaks in similarity and was strongly correlated with prokaryotic community composition. Using a Convergent Cross Mapping network, we identified putative virus-host interactions and six ecological modules, each associated with distinct environmental conditions. The network also revealed putative novel cyanophages with time-lagged correlations to their hosts (late summer) as well as diverse viruses correlated with Nitrososphaerales (winter). By comparison with global metagenomes, we found that 42% of Fram Strait vOTUs peaked in abundance in high latitude regions of both hemispheres (average 61°N and 51°S), and encoded proteins with biochemical signatures of cold adaptation. Our study reveals a rich diversity of polar viruses with pronounced seasonality, providing a foundation for understanding how they regulate and impact ecosystem functionality in changing polar oceans.

## MAIN

Polar regions are subject to the strongest seasonal cycles on Earth, and experience intense pressure from climate change^1,2^. The functioning of polar ecosystems is critical to biogeochemical cycles^3–5^, and is under marked ecological and evolutionary constraints^6^. Such ecosystem processes are strongly driven by microorganisms and their interactions, including viral dynamics. In addition to cell death, viruses also drive evolution via gene exchange, frequency-dependent selection^7,8^, and transmission and expression of auxiliary metabolic genes^9–11^. Therefore, characterizing the diversity and dynamics of viruses is paramount for understanding the function and stability of polar ocean ecosystems.

Relative to more accessible temperate, subtropical, and tropical oceans, few studies have examined virus diversity and ecology in polar waters^12,13^. It is known, however, that polar viromes are distinct from their warm-water counterparts^13–22^ and are typically dominated by bacteriophages with a high level of diversity^14^. In addition to spatial structuring, polar viral communities also shift over time. For instance, total viral counts are typically higher in spring and summer, when constant daylight stimulates phytoplankton growth and the microbial loop^23,24^. Some evidence suggests that different lifestyles of viruses (i.e., lytic vs. lysogenic) may exhibit distinct dynamics across seasons, with lytic infection more prevalent in the Antarctic spring bloom^25^. Furthermore, one study across a calendar year observed seasonal variation among virus communities^22^.

However, due to the challenges of continuous sampling in the polar regions across multiple years, the degree of seasonality among polar viruses – here, meaning annually repeating patterns of populations and communities across the same seasons of different years^26^ – and the potential ecological implications remain to be discovered. In the Arctic, time-series have been critical to advancing an understanding of the Arctic biological carbon pump^27^, elucidating benthopelagic coupling and biotic interactions^28^, both in the water column^29^ and on sinking particles^30^. Furthermore, continuous sampling can discern the impact of ‘Arctic Atlantification’^31–33^: the northward expansion of subarctic habitats through the Fram Strait — the major connection between Atlantic and Arctic Oceans^34^. Here, polar water outflowing the central Arctic Ocean via the East Greenland Current (EGC) meets the West Spitsbergen Current (WSC), transporting warmer Atlantic water into the Arctic Ocean^35–37^. In the WSC, prokaryotic communities exhibit pronounced seasonality, underpinned by changes in photosynthetically active radiation and mixed layer depth^38^. Given this, and the tight coupling of viruses and their hosts^9–11^, we hypothesize that viral populations and communities are seasonally structured in polar regions.

To address this, we examined the diversity and dynamics of viral communities in the Arctic Fram Strait across all seasons over four complete annual cycles at roughly monthly resolution. Our results reveal the diversity and seasonality of Arctic Ocean viruses, their association with environmental conditions and potential microbial hosts, and their distribution across the global ocean.

## RESULTS

We examined 47 long-read metagenomes from samples collected at near-monthly resolution over a four-year period (Aug 2016 – Aug 2020) in the WSC (Fig. 1a). As samples originate from the cellular size-fraction (>0.2 µm), our study characterizes the diversity of actively infecting (i.e. intracellular) viruses, ‘free-living’ viruses with a size of >0.2 µm, viruses attached to particles, and/or integrated viruses.

**Fig. 1.**
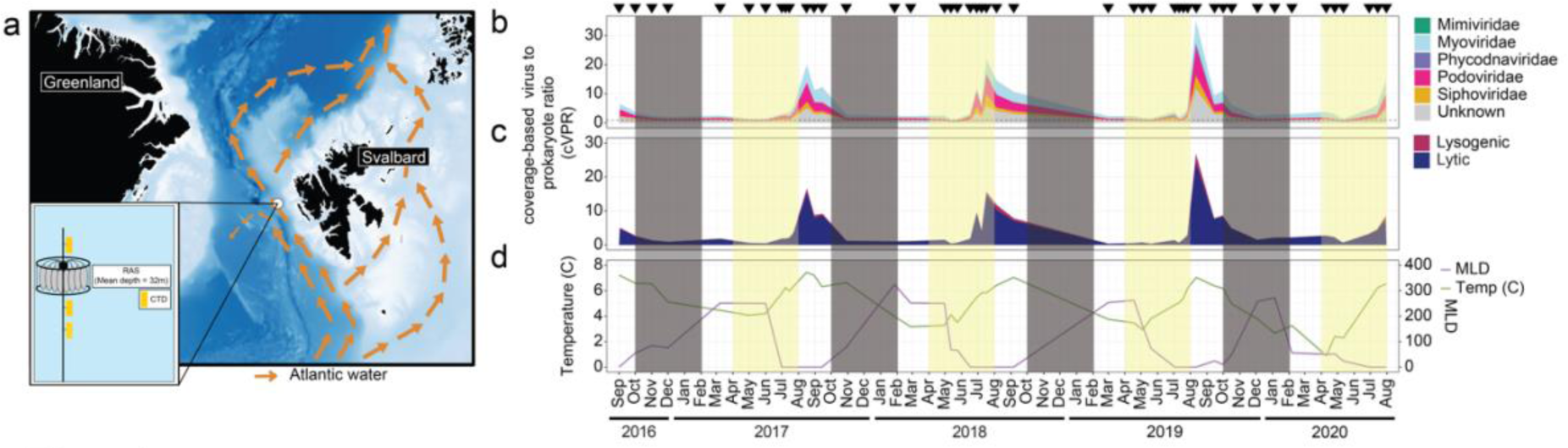
Overview of study site, environmental conditions and viral distributions. **a**, The mooring site in the West Spitsbergen Current of Fram Strait. **b**, cVPR of viral communities across major families detected and **c**, lifestyles during the light and dark cycles from September 2016 to July 2020. **d,** Dynamics of mixed layer depth and temperature which are major community structuring factors. Triangles (top) indicate sampling points.

The total number of reads per sample were 196,489 ± 18,358, with an average length of 5,435 ± 405 bp. Viral sequences were predicted on both assembled contigs and raw reads using a combination of VirSorter2^39^, CheckV^40^, and DeepMicroClass^41^. The overall number of predicted viral sequences was concordant between the two approaches (Supplementary Fig. 1). Hereafter, we focus on the contig-based predictions because of their improved genomic context.

### Diversity of Fram Strait viruses over time

We first investigated how the community of Fram Strait viruses is structured over time. Through contig-based predictions, we identified 5,662 vOTUs (viral Operational Taxonomic Units; 95% sequence clusters, see Methods) that were greater than 10 Kb in length and represented five families. The largest diversity was associated with major bacteriophage groups including 1,990, 1,398, and 506 vOTUs from Myoviridae, Podoviridae, Siphoviridae, respectively. We also detected 81 Phycodnaviridae and seven Mimiviridae vOTUs, but herein focus primarily on prokaryote-infecting viruses.

To assess the temporal dynamics of vOTUs, we normalized their abundance based on the estimated number of prokaryotic genomes in each metagenome - a metric we term coverage-based Virus to Prokaryote Ratio (cVPR). The cVPR approach not only accounts for differences in sequencing depth but also captures shifts in virus to prokaryote-host ratio across samples. Using this metric, we observed clear seasonal structuring in the cVPR of viral communities and taxa. Community cVPR values reached annual maxima of 20-35 and were four-fold higher during July–September (average cVPR 8.6) than during October–June (Fig. 1b). This seasonal variation in cVPR was generally consistent across the major bacteriophage groups. The only seasonal difference at the family level was in Phycodnaviridae, which were more prevalent in late summer (July–September) (Fig. 1b, Supplementary Fig. 1a), when eukaryotic phytoplankton, their presumed hosts, are most abundant ^42^. However, Phycodnaviridae were rare, outnumbered by Myoviridae, Podoviridae, and Siphoviridae bacteriophages by 631-fold, 477-fold, and 154-fold in summer, respectively (Fig. 1b). Overall, lytic lifestyle (as opposed to lysogenic) was predicted to be predominant, with an average of 93.8 ± 0.73% of viruses, a pattern that was invariable across seasons (Fig. 1c), which is in contrast to higher rates of lysogeny in the southern ocean during periods of low production^25^. Thus, some lysogens may be missed by our computational approach or regional differences exist, calling for further investigation into this process using both computational and experimental approaches.

We next investigated how the overall diversity of viruses changes across seasons. To do so, we employed a rarefaction-style strategy that accounts for the high variation in total viruses per sample as well as metagenomes of different sequencing depths. By iteratively rarefying viral counts, we show that the slope of accumulation curves are higher in January-May than June-September (Fig. 2). That is, as sequencing depth increases, viral richness and Shannon diversity increase more rapidly in winter than in summer samples (Fig. 2, Supplementary Fig. 2a). On the other hand, evenness is more similar across months, but lower during May, which corresponds to the timing of phytoplankton bloom conditions (Supplementary Fig. 2b)^43^. Together with the cVPR results, these findings suggest that virus diversity is highest in the winter, while their overall abundance is higher in summer.

**Fig. 2.**
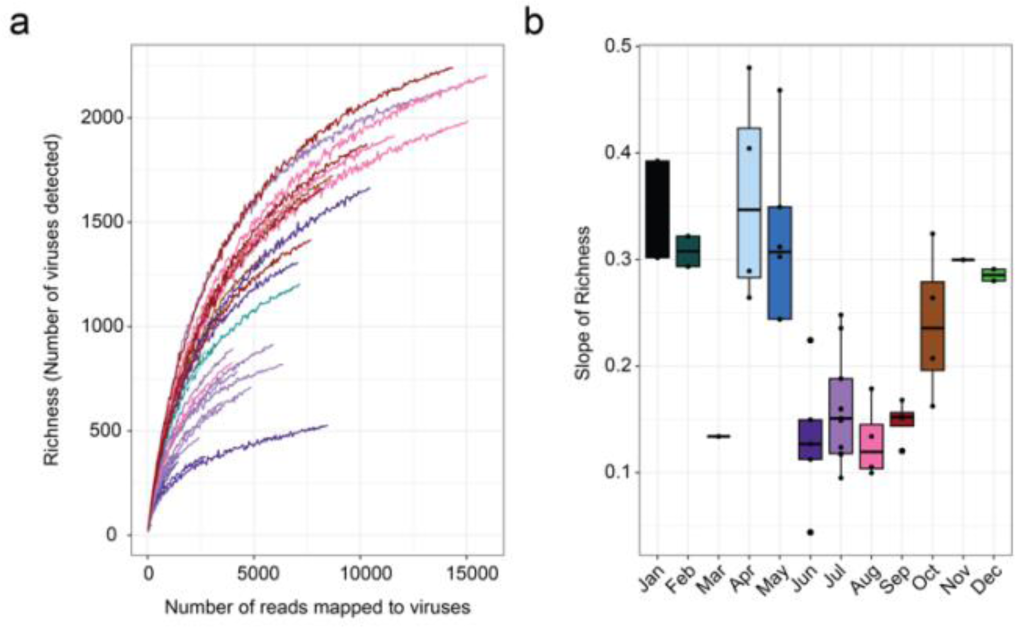
Viral richness seasonally varies. **a**, To account for differences in sequencing depth, vOTU mapping data were subsampled from 25 up to 16,000 viral read counts at 50 count intervals and determined the mean richness at each interval from 100 iterations. The mean richness across subsampled intervals was visualized in a rarefaction-style curve. **b**, Slope of the richness rarefaction curves across sampling months.

In addition to the variations in overall relative abundance (cVPR) and diversity, we also observed strong seasonality in the composition of the major bacteriophage groups (Fig. 3a). The composition of vOTUs within the Myoviridae, Podoviridae, and Siphoviridae showed a sinusoidal-like pattern over time, with peaks in Bray-Curtis similarity at 12, 24, and 36 month intervals (Fig. 3a). Similar observations were made for eukaryotic viruses (families Phycodnaviridae and Mimiviridae), with peaks in similarity at 12-, 24- and 36-month intervals. Furthermore, in some cases, similarity between samples from opposing seasons was zero, which is likely a combination of strong seasonality, low relative abundance (and hence detection) of vOTUs in winter, as well as variable sequencing depth.

**Fig. 3.**
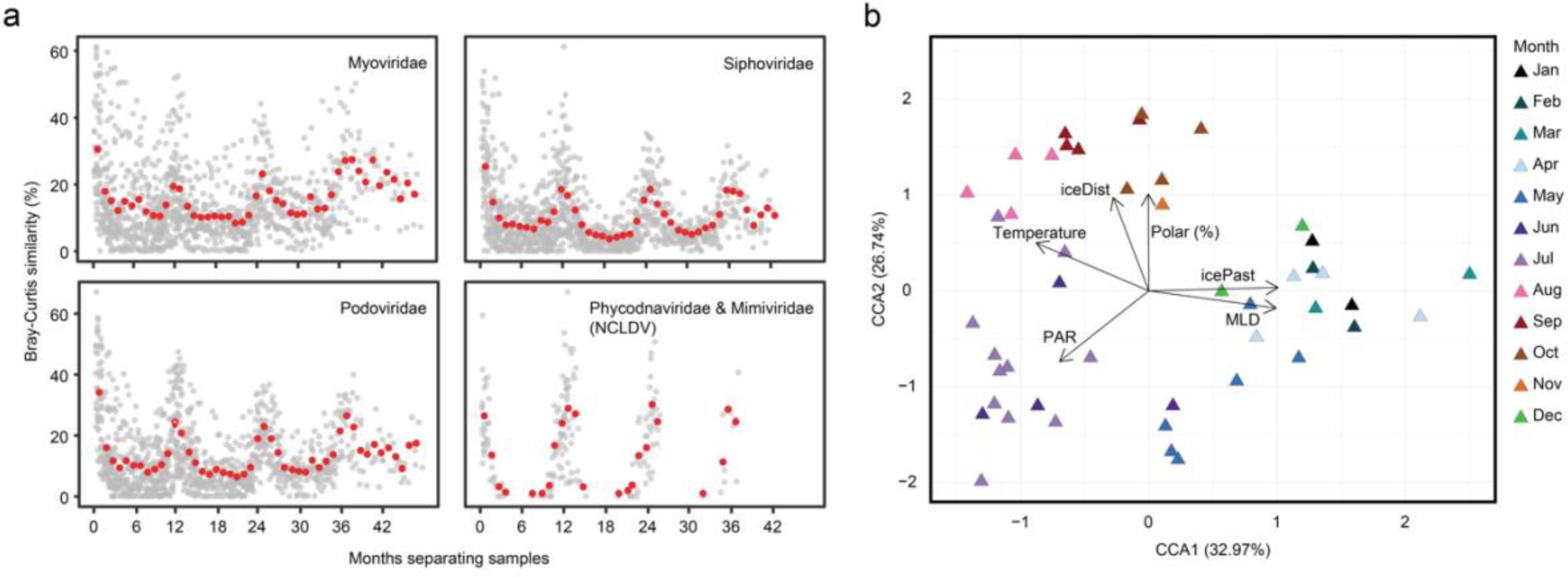
Seasonality of viral communities and their association with environmental conditions. **a**, Bray-Curtis similarity of viral communities over time across four different viral groups: Myoviridae, Siphoviridae, Podoviridae, and NCDLV (Phycodnaviridae and Myoviridae). Each point represents the time between two individual sampling points (x-axis) and their similarity (y-axis) for all pairs of samples. The red points indicate the average similarity for 30.5-day (∼monthly) intervals. **b**, CCA analysis of viral community composition colored by month with vectors representing environmental conditions.

### Virus-host and environmental relationships

To explore the temporal structuring of viruses and their association with prokaryotic hosts, we employed community- and taxon-level analyses in the context of eight physicochemical parameters and prokaryotic community composition data.

At the whole community level, Mantel tests demonstrated the strongest correlation to prokaryotic community composition followed by oxygen and mixed layer depth (MLD) (<u>Supplementary Data 1</u>). To further elucidate physicochemical drivers, we used canonical correspondence analysis (CCA) that revealed a seasonal clustering of samples, with temperature (summer), MLD (winter), PAR (late spring-early summer), and polar water fraction (late summer) accounting for 18.4% of the total variance (Fig. 3b). These results suggest that viral communities are primarily shaped by their hosts, which are in turn driven by environmental conditions plus biological interactions – leading to a complex network of interdependencies and physicochemical linkages, similar to dynamics in temperate environments^7,44^.

Given the coupling between viral and prokaryotic communities, we next explored potential virus-host associations over time at the individual vOTU and 16S rRNA gene amplicon sequence variant (ASV) level. We constructed a Convergent Cross Mapping (CCM) network based on co-occurrences, which includes information about the direction of associations (causal relations) between vOTUs and ASVs. The CCM network comprised 5,136 vOTUs and 850 prokaryotic ASVs with a total of 15,930 directional associations where vOTUs dynamics are ‘following’, or are ‘caused’ by, prokaryotic ASV dynamics (akin to Lotka-Volterra dynamics^45^ (Fig. 4a)). Louvain clustering resulted in six distinct modules, comprising vOTU-ASV associations occurring during the same temporal period. We named the six modules based on their seasonal abundance patterns (Fig. 4b); for example, M1 peaks in late spring, followed by M2. The temporal distinction between modules was confirmed by their correlations with specific physicochemical conditions (Fig. 4c). M1 showed the strongest correlation with daylight, while M2 also had a positive correlation with daylight but a stronger correlation with temperature (Fig. 4c). M3 and M4 peaked in late summer, with M3 having the strongest positive correlation with polar water fraction, and M4 being more positively correlated with temperature. M5 and M6 both negatively correlated with temperature, cVPR, and positively with MLD. However, among these two, only M5 negatively correlated with PAR, being particularly abundant during winter (December-April), while M6 was persistent throughout the year. Generally, the number of vOTUs in a given module outnumbered the number of prokaryotic ASVs, except M5 where ASVs outnumbered vOTUs (Fig. 4d). The overall trend of more vOTUs than ASVs in modules corresponds to the overall larger number of vOTUs vs prokaryotic ASVs examined.

**Fig. 4.**
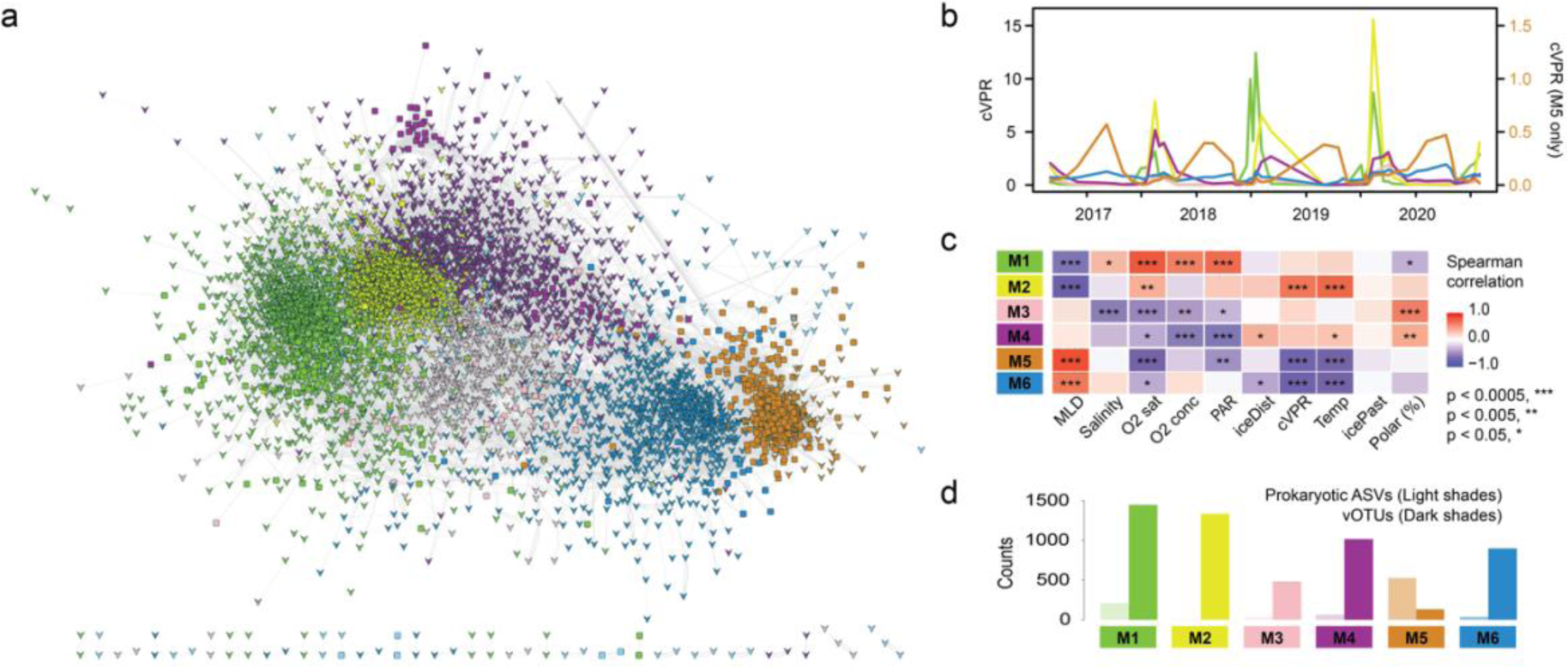
Viral modules and their association with environmental conditions. **a**, Convergent cross mapping network coloured by module. Triangles indicate vOTUs and squares are bacterial ASVs. **b**, Sum of all vOTUs per module over time; note separate y-axis for M5. **c**, Spearman correlation coefficients of modules to environmental parameters, with p-values represented by asterisks. **d**, Count of prokaryotic and viral members of each major module.

The modules were consistently dominated by Myoviridae and Podoviridae (Supplementary Fig. 3a), and the proportion of lytic to lysogenic viruses was rather invariable (Supplementary Fig. 3b). Phycodnaviridae were primarily present in the spring and early summer modules M1 and M2 (Supplementary Fig. 3a) where they constituted 4.3% and 1.2% of the vOTUs, respectively. Together, the major differences between modules, especially relating to bacteriophages, are hence at the vOTU level.

In terms of prokaryotic membership, the modules varied considerably. M1 was dominated by taxa typically associated with copiotrophic conditions or phytoplankton blooms, including Flavobacteriaceae, Rhodobacteraceae, and Porticoccaceae (Supplementary Fig. 3c). In contrast, M5, which contained the largest number of ASVs, comprised diverse prokaryotic taxa, including Pelagibacteraceae, Nitrosopumiliaceae, GCA−002718135 (aka HIMB59), Nitrospinaceae, SAR86, Pirellulaceae, and Planctomycetaceae (Supplementary Fig. 3c). The smaller modules M2, M3, M4 and M6 comprised a variety of taxa, with the most prevalent taxon within each being UBA1611 (Marinimicrobia), and Pelagibacteraceae, respectively (Supplementary Fig. 3c).

In order to evaluate host association patterns, we utilized both the CCM network correlations and host predictions via iPHoP^46^. In total, 22.2% of vOTUs received a host prediction by iPHoP. Those with predicted hosts in the Flavobacteriaceae were predominantly in M1 and M2, while those with predicted hosts in Pelagibacteraceae were more prominent in M3, M4, M5, and M6 (Supplementary Fig. 3d). Among the vOTUs correlated with an ASV in the network, similarly, the most commonly predicted hosts were Flavobacteriaceae, Pelagibacteraceae, Rhodobacteraceae, and Porticoccaceae. However for individual vOTUs and ASVs, only 4.3% of the predictions overlapped at the family level between CCM network correlations and iPHoP. Among these, again, Flavobacteriaceae and Pelagibacteriaceae were the most commonly overlapping, followed by UBA1096 (Pedosphaerales, Verrucomicrobiae) and D2472 (SAR86, Gammaproteobacteria). Overall, the low number of matching CCM correlations and host predictions at the vOTU-ASV level demonstrates the challenge to discern host-virus relationships in diverse and complex ecosystems. Nonetheless, the results provide valuable indications for particular lineages at an overall module level.

### Diversity and dynamics of cyanophages and Nitrososphaerales-associated viruses

From the predicted interactions between vOTUs and ASVs, we further explored the diversity and dynamics of putative cyanophages – as Cyanobacteria are of particular interest with respect to changing conditions in Arctic ecosystems^5,47,48^. We identified putative cyanophage vOTUs based on phylogenetic analysis of *psbA* genes, classification based on VPF-Class^49^, and host prediction via iPHoP^46^. The phylogenetic reconstruction of *psbA* revealed that putative cyanophages are distinct from cultivated relatives from more temperate locations (Fig. 5a). The putative cyanophage vOTUs represented some of the largest viral contigs recovered, with half of the assembled cyanophages being medium-to high-quality, and a quarter >80% complete. Although cyanophage vOTUs all shared *psbA*, their gene content was highly variable between the different families (Cyano-Podoviridae and Cyano-Myoviridae); while within closely related viruses, synteny was usually maintained (Fig. 5b). The cyanophage vOTU abundance peaked during August-September 2017 (Fig. 5c), complementing previous observations of *Synechococcus* in the WSC (Paulsen et al., 2016; Priest et al., 2024). Although the seasonal dynamics were consistent across the cyanophage vOTUs, their maximal cVPR varied from 0.003 to 0.06 (Fig. 5c). In addition, all of the cyanophage vOTUs exhibited lower abundances in 2018-2020 compared to 2017, indicating interannual variation of these viruses and their presumed hosts (Fig. 5c). Overall, the abundance of the cyanophage vOTUs was correlated with the abundance of *Synechococcus* ASVs when considering no time-lag (Spearman’s r = 0.48, p = 0.0008), but a stronger correlation (Spearman’s r = 0.51, p = 0.0008) occurred with a vOTU time-lag of one time-point (average interval of 31.7 days), thus putative cyanophages increased in abundance ∼1 month after *Synechococcus*. For individual ASVs and vOTUs, the strongest correlations were without time delay (n=53), followed by one (n=44) and two (n=32) time-points, indicating some temporal variability between individual cyanophage vOTUs and their putative hosts compared to the groups at-large (<u>Supplementary Data 2</u>).

**Fig. 5.**
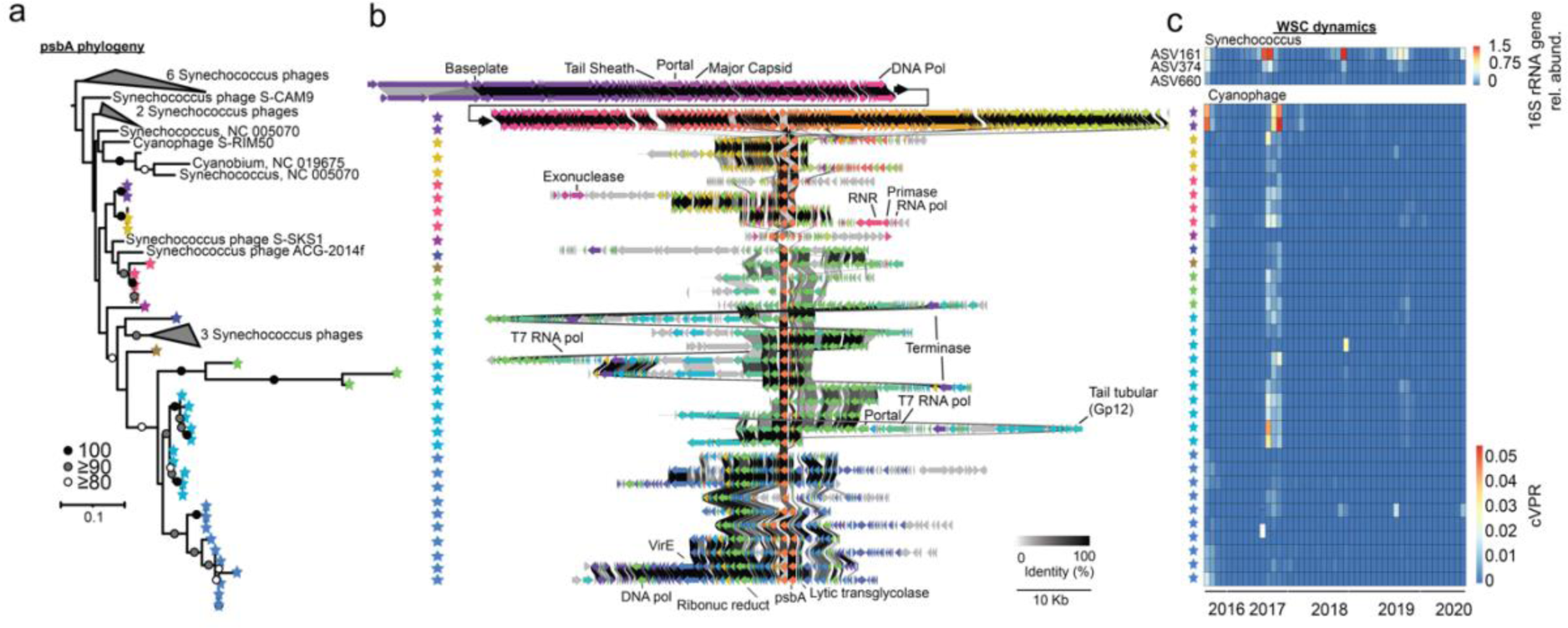
Cyanophages. **a**, *psbA* gene phylogeny of cyanophage vOTUs and reference genomes. **b**, Structure of cyanophage genomes. Coloured stars correspond to phylogenetic groups in (a). The color of the arrows are the same for each ortholog family as defined by sequence similarity by clinker. Bootstrap support is as indicated in figure. percent identity between vOTUs are shown as gray connections as indicated in legend. A 23-gene segment was removed from the longest vOTU contig as indicated by an asterisks for visualization purposes. The region mainly consisted of genes lacking shared orthologous proteins with other viruses. **c**, Dynamics of correlated *Synechococcus* ASVs (top) and cyanophage vOTUS (bottom).

Given the scarcity of data on viruses during the polar night, we also examined the microbial dynamics of viruses related to taxa dominant in winter. To do so, we focused on viruses associated with Nitrososphaerales due to its importance for wintertime nitrogen and carbon cycling ^50,51^ as well as high prevalence in the dataset with the top five Nitrososphaerales ASVs peaking from October-May (average 4.2%) vs. lower abundances from June-September (average 0.47%). We identified associations between 58 vOTUs and five Nitrososphaerales ASVs (Fig. 6), which were primarily associated with the winter module M5. The persistence of these vOTUs despite low virus cVPR demonstrates our ability to identify pronounced seasonality even amongst low abundance viruses. However, more focused investigations on the temporal linkages of these vOTUs and their potential hosts are needed to better understand the environmental impacts of such associations.

**Fig. 6.**
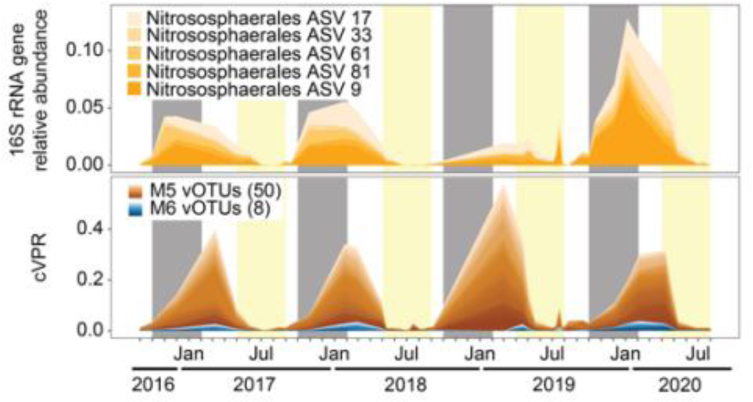
Dynamics of Nitrososphaerales and associated viruses. The shown Nitrososphaerales are the most relatively abundant ASVs, which all are members of M5. Different ASVs and vOTUs are shown by different shades of orange and blue, corresponding to M5 and M6 colors, respectively. The shown vOTUs are those that are correlated with Nitrososphaerales (p < 0.05) in the cross-convergence mapping network (see Methods, Supplementary Fig. 5).

### Bipolarity of Fram Strait viruses

Considering the long-standing discussion on latitudinal microbial diversity gradients^52–55^ and the endemicity of Arctic and Antarctic microbiomes^56,57^, we assessed the distribution of Fram Strait viruses across the global oceans through metagenomic datasets from various large-scale sampling campaigns, such as Malaspina, Tara Oceans and Bio-GO-SHIP (Supplementary Data 3).

Overall, the abundance of Fram Strait viruses peaked in surface seawater (< 200 m) around 60–70° N, with decreasing abundances towards 50° N, and typically no detection in subtropical and tropical surface waters (Fig. 7a). A similar pattern occurred in southern hemisphere surface waters, with peaks in abundance around 45–55° S (Fig. 7a). At individual vOTU level, 42% of vOTUs displayed bi-modal peaks in abundance with separate maxima in each hemisphere (i.e., the two highest peaks in abundance occurred in separate hemispheres). The average peak latitude of vOTUs was 61°N 51°S, respectively (Fig. 7b). Fram Strait viruses were also notably more common in the deep ocean; detected in 87% (338 of 390) of deep global samples, however still more prevalent in northern and southern higher latitudes (Fig. 7b).

**Fig. 7.**
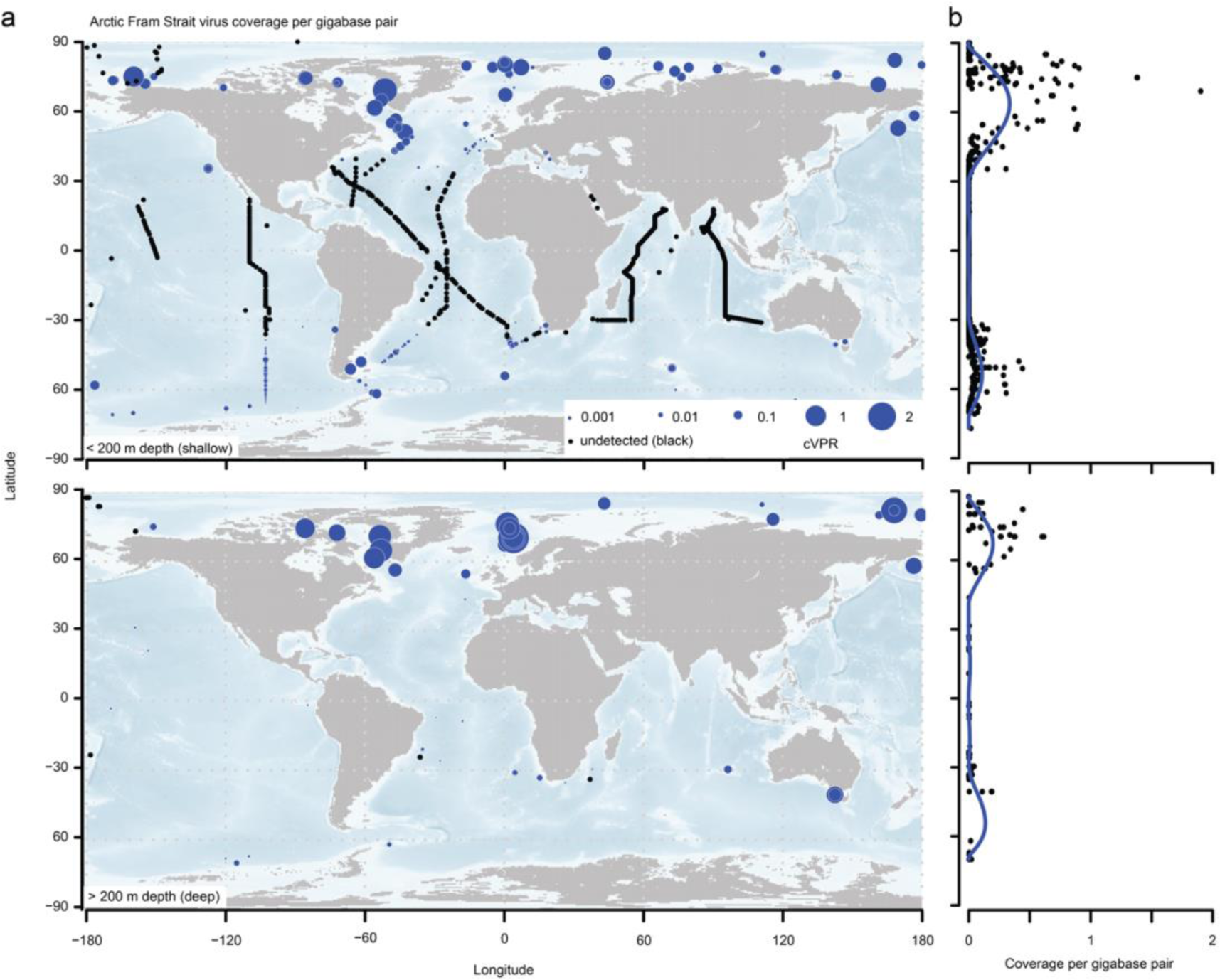
Global distribution of Fram Strait vOTUs by comparison with short-read (0.2–3 μm size-fraction) metagenomes (<u>Supplementary Data 3</u>). **a**, Upper map corresponds to samples collected within the upper 200m and the lower map corresponds to samples collected deeper than 200m. Abundance is calculated as coverage per gigabase pair of sequence. In the ‘deep’ map, latitude and longitude were ‘jittered’ to allow visualization of multiple depths at the same location. **b,** cVPR plotted by latitude across all samples along with a Generalized Additive Model prediction, illustrating the trend of total cVPR by latitude. For more information on samples, see <u>Supplementary Data 3</u>.

Expanding on these observations, we investigated the distributional patterns of the six Fram Strait modules on the global scale. All modules were more prevalent in the northern hemisphere than in the southern hemisphere, but their relative distributions varied, with M1 and M2 being the most abundant in northern surface waters, while M6 being more prominent in higher southern latitudes (Supplementary Fig. 4a), indicating a relatively restricted distribution of M1 and M2. In deeper waters, all modules were less prevalent than in surface waters, with higher prevalence in the northern hemisphere. In deep waters, M6 was the most prevalent (Supplementary Fig. 4b).

### Distinctive amino acid signatures of polar viruses

To expand on the global perspective, we assessed potential adaptations of viruses to polar waters by examining properties that have been attributed to cold adaptation in prokaryotes, in particular amino acid signatures that increase protein flexibility in cold environments. To do this, we examined the amino acid features of proteins from the Fram Strait viruses and from the GOV2.0 dataset, which spans both polar and non-polar sampling locations ^14^. To ensure fair comparisons between the datasets, we focused only on GOV samples originating from ≤35 m water depth (i.e. the average depth of sampling). We found that virus proteins from GOV2.0 from polar waters and Fram Strait were similar in their cold adaptation traits (clustering together), each correlated with latitude, oxygen and temperature (Fig. 8a). Linking the examined protein features with the available environmental data, revealed that the aliphatic index and the nitrogen usage score were positively correlated with temperature (Fig. 8b), whereas other traits of potential cold adaptation, in particular polar charged and uncharged amino acids, showed significant correlation with one or more of the other environmental traits, but not directly with temperature. Together, this suggests that there are biochemical traits distinguishing the polar viruses from the tropical and sub-tropical counterparts encoded in their amino acids.

**Fig. 8.**
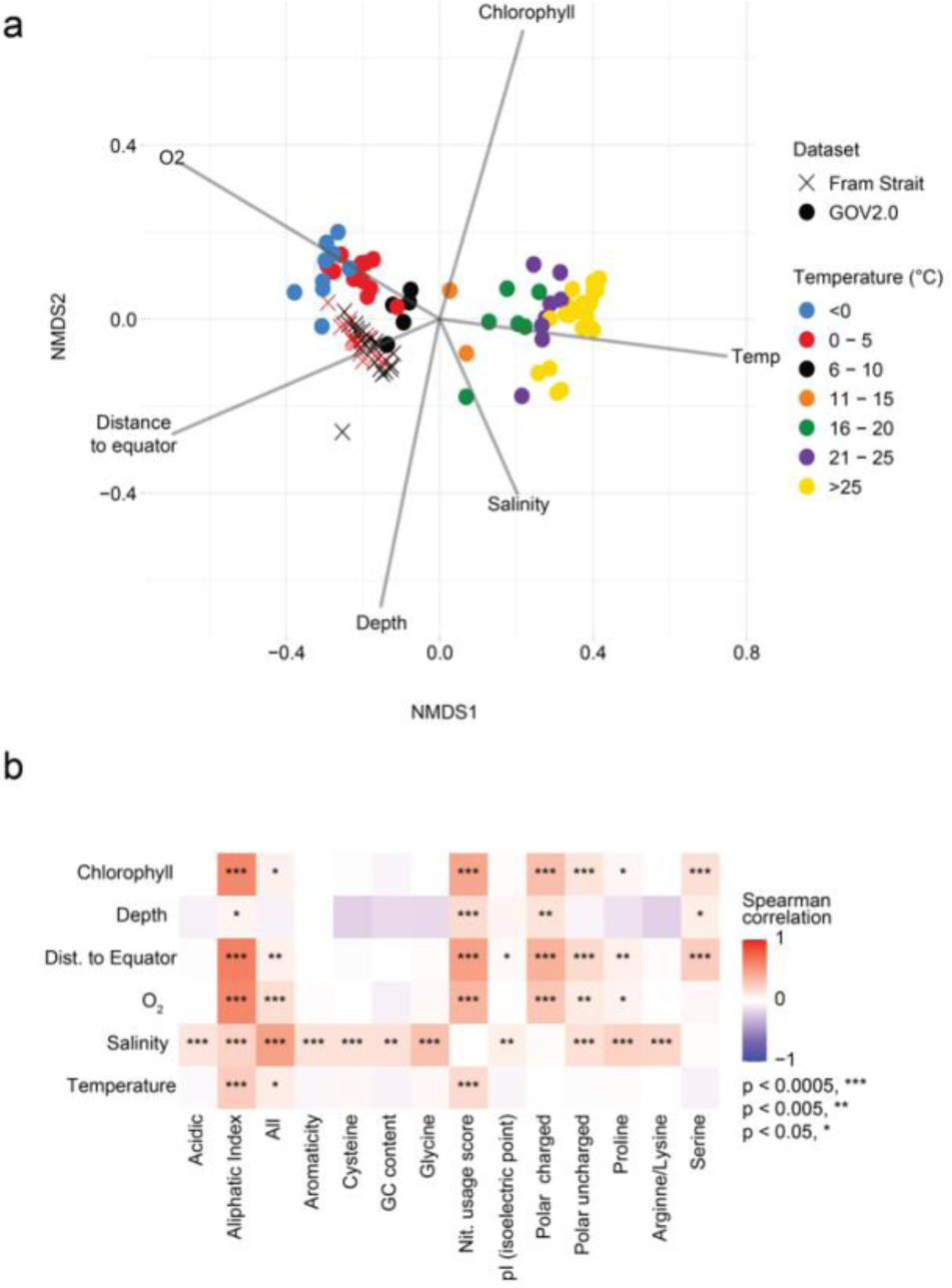
Viral amino acid traits across environmental gradients. **a**, NMDS illustrating the connection between samples and various environmental conditions. **b**, Spearman correlation coefficients of environmental parameters to amino acid traits, with p-values represented by asterisks as indicated.

## DISCUSSION

Harnessing long-read metagenomic data in high temporal resolution, we demonstrate a pronounced, annually repeating seasonality of viruses across multiple years in the Arctic Ocean. The seasonal viral dynamics correlated strongly with both the predominant prokaryotic host communities, as well as environmental parameters. We found that Fram Strait viruses and seasonal modules occur across cold waters of both hemispheres, while being mostly absent from subtropical and tropical waters.

In terms of total virus prevalence, we found strong increases in the ratio of viruses to prokaryotes in the late summer months. The high coverage-based viral abundance estimates in summer are consistent with the timing of increased abundances of free viral particles counted via microscopy (e.g.,^22^), but extend them also to the cellular size-fraction. Our results suggest that the number of viruses in the cellular size-fraction is in the same range (order of magnitude) as that of ‘free’ viruses in seawater, and complement similar metagenomic marker-gene derived ratios of virus-to-prokaryote ratios which have focused so far on size-fractions that include ‘free’ viruses^58^. This observation is consistent with estimated infection rates^10,59^, especially considering that each infected cell can harbor a large number of viral genomes^60–63^.

At the virus community level, the long-read and cellular size-fraction metagenomic approach likely aided assembly of the dominant viruses^64–66^, which often suffer from assembly problems when short-reads are utilized. However, we observed low similarity in opposing seasons compared to studies from more temperate areas^67,68^. The low similarity, reflective of low viral persistence in the ecosystem, may be due to the focus on host-associated viruses, thus excluding free-viruses with no prevalent host that could increase persistence (viral ‘seed-bank’ hypothesis)^69,70^. Furthermore, the low persistence may also be in part related to the low frequency of predicted lysogenic viruses that we detected, which would otherwise be a potential mechanism for viruses with rare hosts to persist^71^. Finally, very low similarity, could be the relatively low coverage overall due to the long-read technology as well as ‘dilution’ of reads from prokaryotes. Nevertheless, the strong seasonality reflects the strongly seasonal host communities ^29,43^.

We observe a pronounced bi-modality in the latitudinal distribution of Fram Strait viruses, with peaks at high latitudes in both the southern and northern hemispheres. In particular, the peaks occur around the Northern and Southern Polar fronts (70°N / 60°S)^72–75^, suggesting that these boundaries of oceanic realms represent a hotspot of polar-adapted viruses, possibly due to the pronounced biological and physicochemical gradients that are present, and overall greater productivity. It might be, however, somewhat biased by a larger number of samples collected along these fronts. The association to high-latitudes of the Arctic viruses is also reflected within their amino acid sequences, with signatures of cold adaptation that may contribute to their success in colder waters, thus expanding previous observations from the southern ocean^18^ and polar eukaryotic viruses^76^). Further investigations to unravel the diversity and dynamics of viruses in the two polar regions are crucial, given that climate shifts are causing major biological and physicochemical perturbations in these regions.

The peaks of Arctic viruses during summer and their specialization to high-latitude regions, in general, call for further process and targeted studies to elucidate the host communities that they interact with and influence. Previously, viruses of phytoplankton (cyanobacteria and eukaryotic phytoplankton) have been experimentally observed to exert less top-down pressure than eukaryotic grazers at high latitudes^77,78^, but how this relates to the total microbial community remains unknown. In any case, our study demonstrates that the impacts of Arctic viruses are likely very different depending on season and ecosystem state.

In conclusion, our study advances the basis for understanding how viruses regulate and impact the dynamic and changing polar ecosystem, setting the stage for more detailed population dynamics studies, as well as process- and host-specific studies. Extended time-series observations will allow for improved understanding of the Arctic ecosystem impacts, in this ecosystem undergoing rapid change.

## METHODS

### Seawater collection and eDNA sequencing

Moored Remote Access Samplers (RAS; McLane) autonomously collected and fixed seawater in the eastern and western Fram Strait (Fig. 1) at weekly to fortnightly intervals between 2016–2020^38,51^. Sampling occurred in the framework of the FRAM / HAUSGARTEN Observatory. The resulting eDNA was used to sequence 16S rRNA gene fragments using primers 515F–926R^79^, processed into Amplicon Sequence Variants (ASVs) using DADA2 as described under https://github.com/matthiaswietz/FRAM_eDNA. Subsequently, we only considered ASVs with ≥3 reads in ≥3 samples, corresponding to a total of 3,748 ASVs. DNA extracts from selected timepoints were additionally used to generate PacBio HiFi metagenomes at the Max Planck Genome Centre, Cologne, Germany. Further details about the molecular analyses are described in^38,51^. In total, we herein report 94 amplicon samples and 47 metagenomes from the WSC, and 9 metagenomes from the EGC (<u>Supplementary Data 4</u>).

### Environmental parameters

Attached to the RAS were Seabird SBE37-ODO CTD sensors that measured temperature, depth, salinity, and oxygen concentration. Sensor measurements were averaged over 4 h around each seawater sampling event. Physical sensors were manufacturer-calibrated and processed in accordance with https://epic.awi.de/id/eprint/43137. Employing multiple CTD sensors along the mooring depths enabled the determination of the minimum mixed layer depth (MLD) at each sampling time point. Chlorophyll concentrations were measured via Wetlab Ecotriplet sensors. Surface water Photosynthetically Active Radiation (PAR) data, with a 4 km grid resolution, was obtained from AQUA-MODIS (Level-3 mapped; SeaWiFS, NASA) and extracted in QGIS v3.14.16 (http://www.qgis.org).

### Virus assembly, prediction, classification, and host prediction

Long-reads from each sample were assembled individually using hifiasm-meta^80^ with default settings. Viruses were predicted from both assembled contigs and the long-reads themselves using a combination of VirSorter2^39^ and DeepMicroClass^41^. For VirSorter2, we included all possible viral groups: dsDNAphages, RNA viruses, ssDNA viruses, nucleocytoplasmic large DNA viruses (NCLDV), and Lavidaviridae. CheckV^40^ was then used to quality check the long-reads and contigs identified as viruses by VirSorter2 as well as to trim potential host regions from identified proviruses. Contigs identified as viruses by VirSorter2 (--min-score 0.5) with at least one viral gene (predicted by CheckV) were further screened using DeepMicroClass. Contigs that were classified as either eukaryotic or prokaryotic by DeepMicroClass were discarded. Contigs that were identified as RNA viruses by VirSorter2 were not analyzed. We clustered the remaining viral contigs into vOTUs, as defined previously^81^, using cd-hit with the following parameters: cd-hit-est -M 100000 -c 0.95 -d 100 -g 1 -aS 0.85.

For classification, we used VPF-Class (default settings) which uses sequence similarity of viral proteins to reference virus sequences^49^. VPF-Class utilizes the Baltimore classification system which is based on morphological and replication attributes of viruses. For VPF-Class-based taxonomy, families were assigned when both membership ratio and confidence score were >0.5. Viral contigs were also analyzed using VIBRANT^82^ to differentiate between lytic and lysogenic lifestyles. Although VPF-Class is capable of generating host predictions on its own, we supplemented these predictions by using iPHoP v.1.3.3^46,83–86^ with default settings, along with a custom database by adding previously assembled bacterial and archaeal MAGs from the Fram Strait (PRJEB67368) and removed MAGs from one oceanic study that was not manually curated, due to their high degree of potential virus contamination (PRJNA385857), which could result in false positives. When multiple host predictions were available from the combined iPHoP and RaFAH outputs, we selected the prediction with the highest confidence score.

### Mapping and normalization of viral abundance

Metagenomic long-reads from all samples were mapped to vOTUs (>10 Kb, based on assembly-based and raw-read approaches) using minimap2 (parameter: asm5)^87^ and only the primary alignments were considered in subsequent analyses.

To quantify the abundances of vOTUs across metagenome samples, we employed a two-step normalization process to account for differences in read lengths and sequencing depth. First, we determined the mean coverage across the viral sequence length from mapped read counts (with a threshold that >25% of the viral sequence must be covered). Second, we divided the mean coverage by the estimated number of prokaryotic genomes in each metagenome, as predicted based on the mean coverage across universal single copy genes^88^. We call this metric coverage-based virus to prokaryote ratio (cVPR). This approach is similar to that used to derive virus to microbial ratios on virus to prokaryotic marker genes elsewhere^58^, but considers the full viral coverage. In addition to this, we also calculated the coverage of each virus per gigabase pair (cVGB) of metagenome for a given sample and demonstrated that the two metrics yield comparable values (Supplementary Fig. 1).

### Estimation of viral diversity

Given that the vOTUs were derived from metagenomes with different sequencing depths, we employed an iterative subsampling approach to quantify and compare the alpha diversity of viruses across samples. For this, we applied 100 iterations of subsampling the vOTU count table at a range of different depths, spanning from 25 up to 16,000 counts at 50 count intervals. For every iteration, richness, evenness and Shannon diversity was calculated, with mean values being determined for each 50-count interval. The functions *rrarefy, specnumber* and *diversity* from the vegan package were used to perform subsampling and alpha diversity calculations. The mean values were used to generate a rarefaction-style curve of alpha diversity using the *ggplot2* package. To compare shifts in alpha diversity over time, we performed a linear regression between the mean alpha diversity values and the subsampling depths. From this, we determined and compared the steepness of the slopes across sampling months, which indicate the rate of change in richness, evenness and Shannon diversity with changes in sampling depth (virus counts).

### Co-occurrence network

To identify temporal co-occurrence patterns, cVPR values per vOTU and relative abundance of 16S rRNA gene amplicon ASVs were converted into temporal profiles by Fourier transformation. Temporal profiles were constructed based on 16 Fourier coefficients, which capture the majority of observed vOTU and ASV peaks within the four-year period. Pairwise correlations between individual temporal profiles were then computed between all vOTUs and ASVs. Higher Pearson correlation values indicated similar temporal profiles. For network construction, we first calculated Pearson correlations for all pairs resulting in an undirected graph, from which we only considered correlations >0.7 after multiple testing corrections using the Benjamini-Hochberg procedure. To delineate strongly connected components representing co-occurring taxa, the Louvain community detection algorithm was applied to the entire graph^89^. Next, the putative associations were further evaluated based on Convergent Cross Mapping (CCM) in order to discern causal relationships between taxa in time-series data, as outlined by^90^. CCM enables the prediction of a species’ time-series based on the knowledge of another species’ time-series. Initially, we constructed a CCM network encompassing all pairwise combinations. Subsequently, we extracted the in- and outgoing edges between nodes that were also connected in the co-occurrence network, utilizing resources from https://github.com/rakro101/otter. To quantify the strength of relationships, we employed Normalized Mutual Information (NMI) to account for nonlinear relations^42^. A permutation approach was employed to compute significance values for edge weights, with the objective of determining whether the NMI values exceeded those expected for random edges. Further details on the construction and validation of the CCM network can be found in^42^. The CCM network was visualized in Cytoscape v.3.10.1. Additionally, to identify potential time-lagged correlations, we employed extended local similarity analysis ^91^ with a maximum delay of two sampling points on the original non-Fourier transformed data.

### Statistical analyses

Bray-Curtis similarity, Mantel tests, CCA, and Spearman correlation were performed in R v. 4.3.1 using the *vegan* and *Hmisc* packages. We included only vOTUs with a breadth of coverage > 0.25.

### Cyanophage phylogenetics and comparative genomics

Putative cyanophages were predicted via VPF-class and iPHoP predictions. Subsequently, we focused on the most confident predictions by selecting cyanophages harboring the *psbA* core photosystem, identified by hmmscan of viral predicted proteins (prodigal^92^) PFAM for Photo_RC (PF00124). Viral and prokaryotic reference *psbA* were extracted from the vConTACT2 viral reference database (ViralRefSeq-prokaryotes-v211) and GTDB prokaryotic reference database (version 214^93^) by hmmscan with PFAM Photo_RC. Eukaryotic reference *psbA* genes were extracted via UniRef. A preliminary tree was constructed from these sequences to differentiate between D1 (PsbA) and D2 (PsbD) photosystem II (PSII) reaction centre proteins. To do so, we aligned using MAFFT^94^ followed by trimming with trimAl^95^, and FastTree^96^ for phylogenetic reconstruction. PsbD were identified manually and excluded. The remaining sequences were considered PsbA and phylogenetic reconstruction performed a second time with the same alignment and trimming steps; at this stage, tree building was performed with IQ-TREE^97^. In order to construct synteny plots, the PsbA-containing contigs were annotated with Prokka^98^, and then visualized with clinker^99^.

### Mapping to global metagenomes

Metagenomes from ten oceanic datasets, including Tara Oceans, Malaspina, and Bio-GO-SHIP (<u>Supplementary Data 3</u>), were downloaded from NCBI, and quality-filtered with Trimmomatic (parameters: LEADING:3 TRAILING:3 SLIDINGWINDOW:50:30 MINLEN:50)^100^. Only DNA samples collected from discrete depths (e.g., not composites of the mixed layers or similar) and the prokaryotic size-fraction (0.2–3 µm) were used to allow comparison with our dataset. Reads were mapped to the clustered contigs using bbmap.sh from the BBTools package (version 39.01, parameters: minid=95 idfilt=95). Trimmed mean coverage was calculated via CoverM^101^ on contigs with minimum covered fraction of >25% for each sample. Trimmed mean coverage values were then divided by total Gbp per quality filtered paired reads. For determining abundance trends in the total dataset and at the module level, we computed a Generalized Additive Module (GAM) between cVPR and latitude using the gam function from the *mgcv* package^102^, with ten degrees of freedom (k=10). For plotting the predicted GAM, negative values were converted to 0. For calculating the maximum latitude that viruses occurred in each hemisphere, we used the GAM model, computing where ‘peaks’ occur and then determining which two peaks had the highest abundance.

### Calculation of amino acid traits

Proteins were predicted from GOV2.0 and Fram Strait viruses via Prodigal v.2.6.3 (parameter: -p meta)^92^. Then, amino acid composition and various biochemical properties of the predicted protein sequences were assessed using custom R scripts, which are available via GitHub (https://github.com/alyzzabc/fram_strait_viruses_2016-2020). Briefly, protein sequences containing the ambiguous amino acid “X” were removed, followed by counting the occurrences of each proteinogenic amino acid. The frequencies of single amino acids (G, S, P, C) and of amino acid groups (acidic, polar uncharged, polar charged, aromatic) were calculated relative to the protein length. For the amino acids arginine (R) and lysine (K), the ratio R/K was calculated instead of occurrence. The molecular weight was calculated according to^103^ and for calculation of the aliphatic index we followed^104^. The pI function from the *Peptides* package was used to calculate the pI of the predicted proteins using the EMBOSS scale. The nitrogen usage score (NUS) was calculated using the following formula:

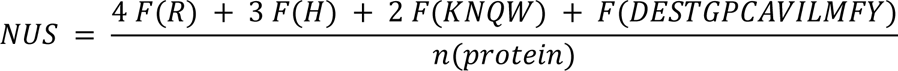

where F(X) corresponds to the absolute count of each amino acid and n(protein) to length of the protein.

To analyze the variation of the viral protein features in relation to the environmental conditions, we filtered out samples collected from a water depth >35 m followed by calculation of the Bray-Curtis distance using the *metaMDS* function from the *vegan* package.

To calculate pairwise correlations, the *vegan* package was used: the distances for each protein feature (Bray-Curtis) and environmental condition (Euclidean) were calculated using the *dist* function. Pairwise Mantel tests were done using the Spearman method with 9999 permutations via the *mantel* function. The NMDS plot and the correlation heatmap were prepared using the *ggplot2* package.

## ACKNOWLEDGEMENTS

Christina Bienhold, Katja Metfies, Ian Salter and Antje Boetius co-designed the mooring strategy, and coordinated sample collection and processing. Katja Metfies coordinated amplicon sequencing. Wilken-Jon von Appen, Sinhué Torres-Valdés, and Daniel Scholz carried out physicochemical and oceanographic measurements. We thank Jana Bäger, Theresa Hargesheimer, Jakob Barz, Anja Batzke, Rafael Stiens, and Lili Hufnagel for RAS operations; Normen Lochthofen, Janine Ludszuweit, Lennard Frommhold and Jonas Hagemann for mooring operations; Jakob Barz, Swantje Ziemann and Anja Batzke for DNA extraction, amplicon library preparation and sequencing; and Bruno Huettel, Christian Woehle and the technicians at the Max Planck Genome Centre in Cologne for metagenome sequencing. Captains, crew and scientists of RV Polarstern cruises PS99.2, PS107, PS114, PS121 and PS126 are gratefully acknowledged. This project has received funding from Polarstern grants AWI_PS99_00, AWI_PS107_05, AWI_PS114_01, AWI_PS121_07, AWI_PS126_05, and AWI_PS126_07. Further support came from the Helmholtz Association, the Max Planck Society, and a Helmholtz Young Investigator Grant to DMN.

## DATA AVAILABILITY

Raw metagenomic reads are available under ENA BioProjects PRJEB67368 (WSC) and PRJEB52171 (EGC). 16S rRNA amplicon reads are available under PRJEB43890 (2016-2017), PRJEB43889 (2017-2018), PRJEB67813 (2018-2019), and PRJEB66202 (2019-2020). Physicochemical parameters are available at PANGAEA under https://doi.pangaea.de/10.1594/PANGAEA.904565 (2016-2017), https://doi.pangaea.de/10.1594/PANGAEA.904534 (2017-2018), https://doi.pangaea.de/10.1594/PANGAEA.941126 (2018-2019), and https://doi.pangaea.de/10.1594/PANGAEA.946508 (2019-2020). Code for reproducing workflows and figures are available via GitHub https://github.com/alyzzabc/fram_strait_viruses_2016-2020/.

## AUTHOR CONTRIBUTIONS

AC identified, annotated, mapped, and assembled viruses. AC and DMN analyzed viral and prokaryotic community data. TP contributed normalization techniques and alpha diversity metrics. EO and OP performed network analysis and module identification. JM performed amino acid analyses. MW and DMN instigated the study. DMN designed and coordinated the analyses. AC and DMN wrote the paper with input from all co-authors, especially TP and MW.

