## Supplementary Figures for "Arctic Ocean virus communities: seasonality, bipolarity, and prokaryotic interactions"

### SUPPLEMENTARY FIGURES AND LEGENDS

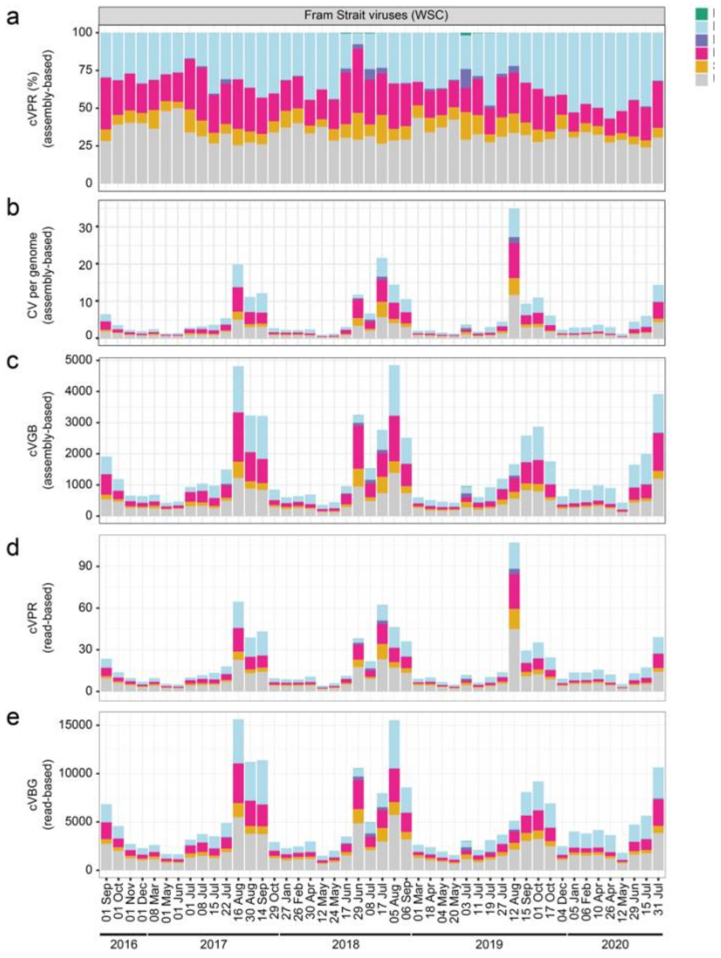


S. Figure 1

**Supplementary Fig. 1**. Normalized abundances of viral families. **a**, Proportion of assembly-based cVPR (coverage based virus:prokaryote ratio). **b**, Total mapping and assembly-based cVPR. **c**, Total mapping and assembly-based coverage per gigabase pair (cVGB). **d**, Total read-based (i.e., non mapping and assembly-based) cVPR. **e**, Total read-based cVGB. The results indicate that the overall patterns are robust across different methods for calculating abundance.

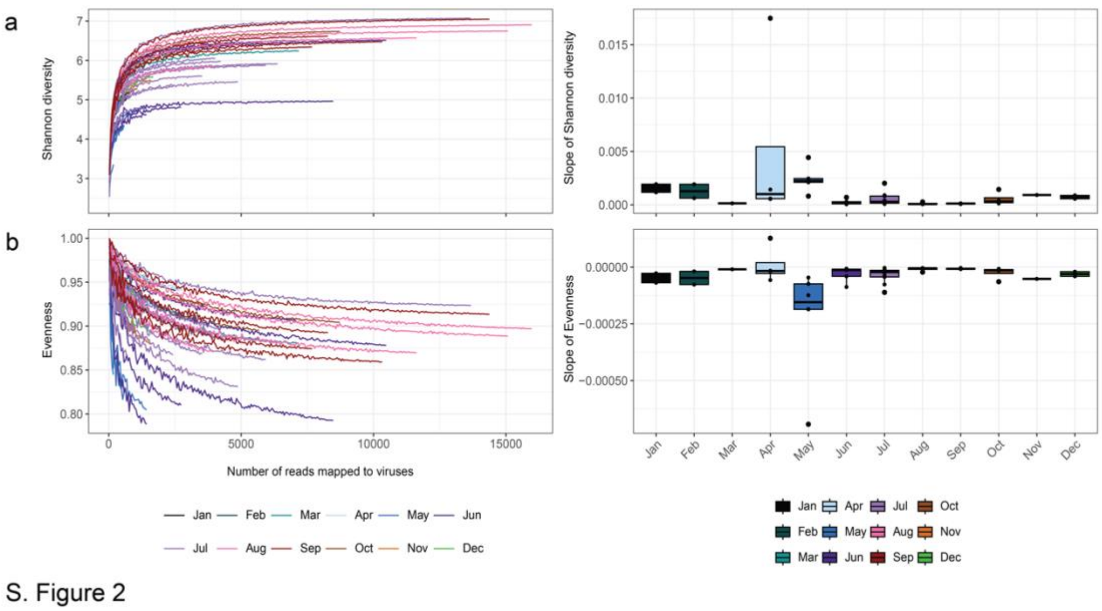


**Supplementary Fig. 2.** Seasonal shifts in Shannon diversity and evenness of vOTUs. **a**, similar to Fig. 2, vOTU mapping data was subsampled spanning from 25 up to 16,000 viral read counts at 50 count intervals and determined the mean Shannon diversity and evenness at each interval from 100 iterations. The mean values were visualized in rarefaction-style curves. **b**, Slopes of the Shannon diversity and evenness rarefaction curves across sampling months.

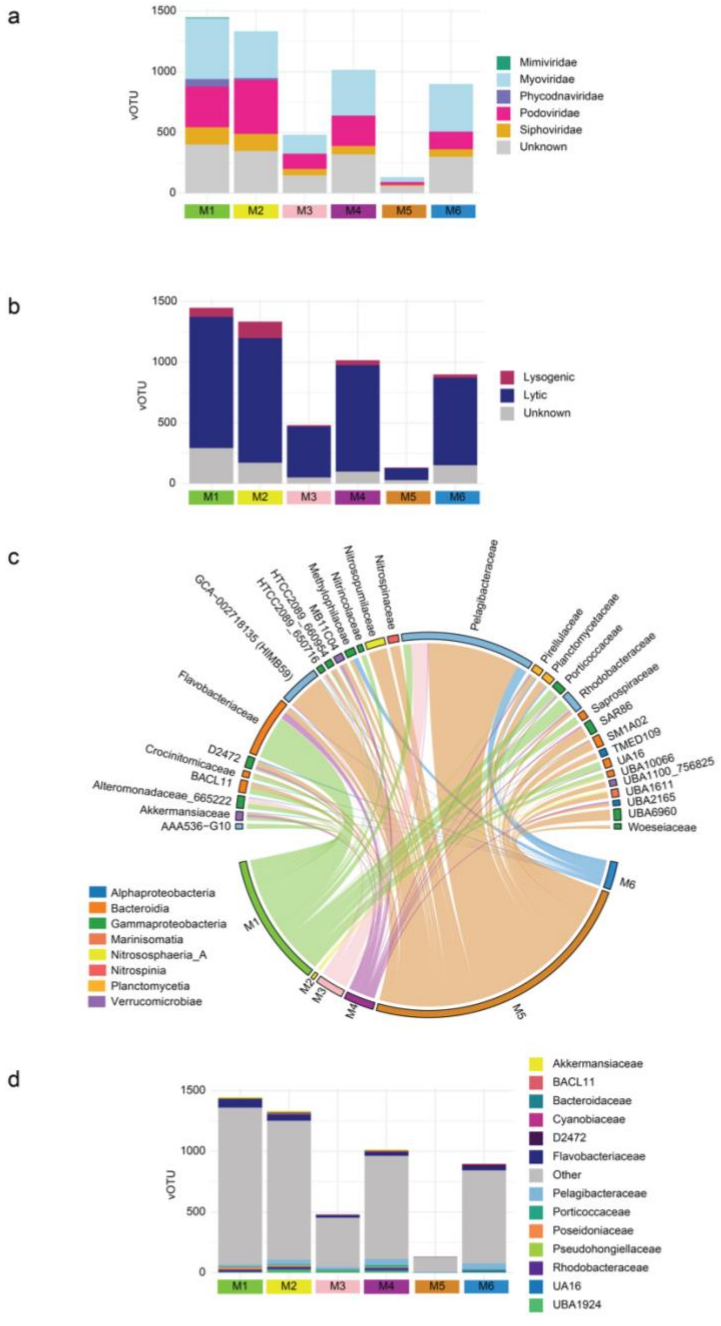

S. Figure 3

**Supplementary Fig. 3**. Overview of the membership and characteristics of taxa within the major Louvain modules. **a**, Family-level classification of vOTUs in the respective modules. **b**, Viral lifestyle prediction based on VIBRANT. **c**, Taxonomy of prokaryotic ASVs by module; only taxa with greater than five occurrences are shown. UBA1096 is excluded because there were only 5 ASVs (all occurring in M5). **d**, iPHoP-predicted hosts of the vOTUs.

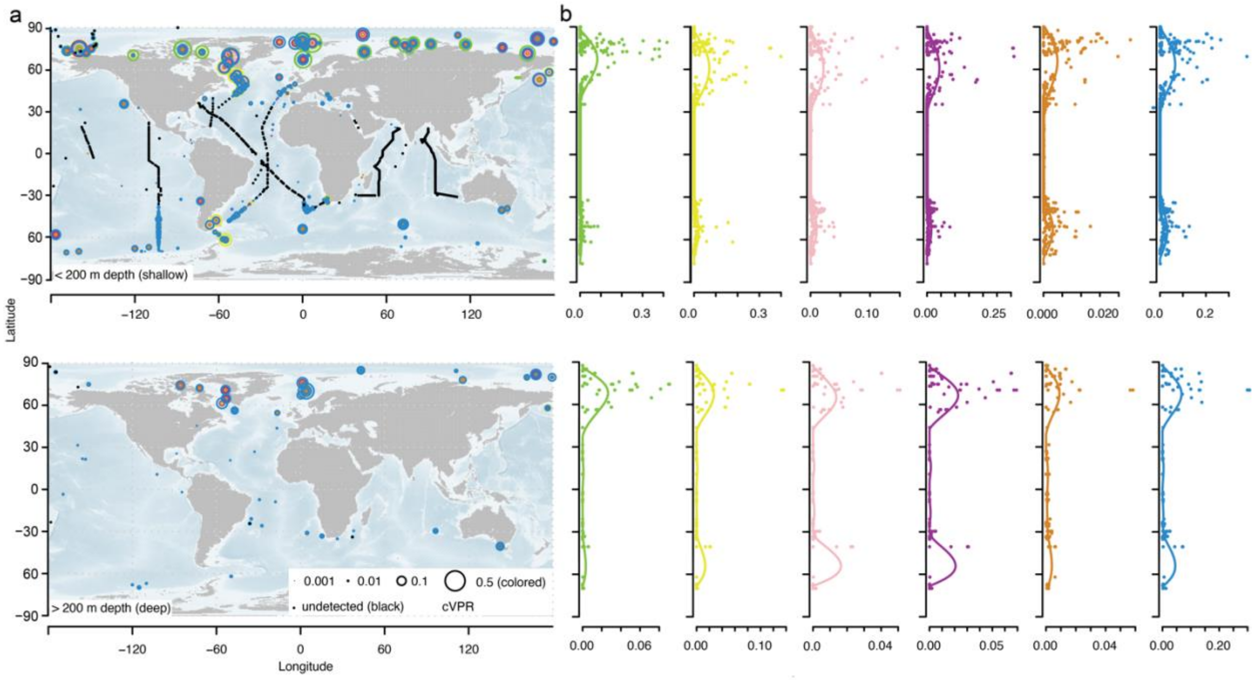


**Supplementary Fig. 4**. Global distribution of Fram Strait vOTUs at the module level. **a**, Similar to Fig. 7, the upper panel is total coverage per gigabase pair for samples collected from less than 200m (surface) and the lower panel is for samples from greater than 200m (deep). **b,** Latitudinal gradient for modules, with a Generalized Additive Model prediction illustrating the trend of the modules across latitude. For more information on samples, see Supplementary Data 3.

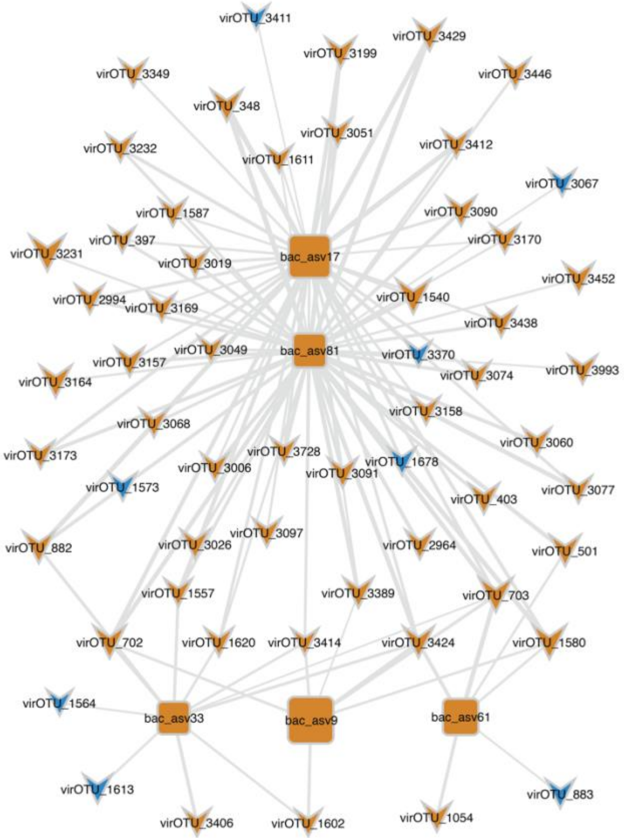


S. Figure 5

**Supplementary Fig. 5**. Convergent cross mapping networking of abundant Nitrososphaerales and vOTUs. Edge weight corresponds to strength of correlation (minimum shown; all $p<0.05$). Node size corresponds to the sum of abundance across the time-series. Node color corresponds to the module membership of the ASV and vOTUs.

#### SUPPLEMENTARY DATA LEGENDS

**Supplementary Data 1.** Mantel r statistics for various combinations of biological and environmental parameters. Statistics were determined using Bray-Curtis distances computed with different filtering criteria: all vOTUs, no NCLDV vOTUs, no unknown vOTUs, and no NCLDV or unknown vOTUs. The additional columns beyond "All vOTUs" were added because, beyond the physicochemical conditions, only prokaryotic community composition was compared (not eukaryotic hosts), so we determined the similarity with prokaryotic community also after excluding eukaryotic hosts. The differences were negligible.

**Supplementary Data 2.** eLSA results for cyanobacteria and putative cyanophages.

**Supplementary Data 3.** Project accession information of public data used in mapping Fram Strait viruses to global metagenomes.

**Supplementary Data 4.** Sample information about Fram Strait metagenomes used in the current study.
